# Neuraminidase antigenic drift of Influenza A virus H3N2 clade 3c.2a viruses alters virus replication, enzymatic activity and inhibitory antibody binding

**DOI:** 10.1101/2020.02.20.957399

**Authors:** Harrison Powell, Andrew Pekosz

## Abstract

In the 2014-2015 influenza season a novel neuraminidase (NA) genotype emerged in the Johns Hopkins Center of Excellence for Influenza Research and Surveillance (JH CEIRS) surveillance network as well as globally. This novel genotype encoded a glycosylation site at position 245-247 in the NA protein from clade 3c.2a H3N2 viruses. In the years following the 2014-2015 season, this novel NA glycosylation genotype quickly dominated the human H3N2 population of viruses. To assess the effect this novel glycosylation has on virus fitness and antibody binding, recombinant viruses with (NA Gly+) or without (NA Gly-) the novel NA glycosylation were created. Viruses with the 245 NA Gly+ genotype grew to a significantly lower infectious virus titer on primary, differentiated human nasal epithelial cells (hNEC) compared to viruses with the 245 NA Gly-genotype, but growth was similar on immortalized cells. The 245 NA Gly+ blocked human and rabbit monoclonal antibodies that target the enzymatic site from binding to their epitope. Additionally, viruses with the 245 NA Gly+ genotype had significantly lower enzymatic activity compared to viruses with the 245 NA Gly-genotype. Human monoclonal antibodies that target residues near the 245 NA glycosylation were less effective at inhibiting NA enzymatic activity and virus replication of viruses encoding an NA Gly+ protein compared to ones encoding NA Gly-protein. Additionally, a recombinant H6N2 virus with the 245 NA Gly+ protein was more resistant to enzymatic inhibition from convalescent serum from H3N2-infected humans compared to viruses with the 245 NA Gly-genotype. Finally, the 245 NA Gly+ protected from NA antibody mediated virus neutralization. These results suggest that while the 245 NA Gly+ decreases virus replication in hNECs and decreases enzymatic activity, the glycosylation blocks the binding of monoclonal and human serum NA specific antibodies that would otherwise inhibit enzymatic activity and virus replication.

**Author Summary:** Influenza virus infects millions of people worldwide and leads to thousands of deaths and millions in economic loss each year. During the 2014/2015 season circulating human H3N2 viruses acquired a novel mutation in the neuraminidase (NA) protein. This mutation has since fixed in human H3N2 viruses. This mutation at position 245 through 247 in the amino acid sequence of NA encoded an N-linked glycosylation. Here, we studied how this N-linked glycosylation impacts virus fitness and protein function. We found that this N-linked glycosylation on the NA protein decreased viral replication fitness on human nasal epithelial cells (hNEC) but not immortalized Madin-Darby Canine Kidney (MDCK) cells. We also determined this glycosylation decreases NA enzymatic activity, enzyme kinetics and affinity for substrate. Furthermore, we show that this N-linked glycosylation at position 245 blocks some NA specific inhibitory antibodies from binding to the protein, inhibiting enzymatic activity, and inhibiting viral replication. Finally, we showed that viruses with the novel 245 N-linked glycosylation are more resistant to convalescent human serum antibody mediated enzymatic inhibition. While this 245 N-linked Glycan decreases viral replication and enzymatic activity, the 245 N-linked glycosylation protects the virus from certain NA specific inhibitory antibodies. Our study provides new insight into the function of this dominant H3N2 NA mutation and how it impacts antigenicity and fitness of circulating H3N2 viruses.

## Introduction

Each year seasonal influenza accounts for 3 to 5 million incidences of severe disease and up to 650,000 deaths [1]. Most influenza vaccines rely on the generation of antibodies against the hemagglutinin (HA) protein, one of the two major glycoproteins on the virion surface. The anti-HA protein antibodies inhibit virus entry into cells but also provide an immune pressure which leads to the emergence of virus strains with mutations in HA antigenic sites [2, 3]. This antigenic drift leads to escape from vaccine- and infection-induced immunity and results in the need to change influenza vaccine strains on a fairly frequent basis.

There is renewed interest in generating influenza vaccines that provide broader and stronger protection against several virus strains [4-6] and the other major influenza surface glycoprotein, the neuraminidase (NA) protein, has emerged as a potential candidate for such a universal influenza vaccine [6]. The NA protein has a neuraminidase activity that is critical in two stages of the virus life cycle[7-9]. The NA protein cleaves sialic acid from mucins that coat airway epithelial cells which reduces HA protein binding to mucins and facilitates entry into respiratory epithelial cells[10]. The neuraminidase activity also removes sialic acid from host cell membrane bound proteins and viral HA and NA proteins at late times post infection, allowing viral particles to efficiently bud and spread to other respiratory epithelial cells[7, 11].

Anti-NA antibodies can prevent or decrease the severity of influenza infection[12-15]. High titer anti-NA antibodies have been correlated with decrease disease severity and protection in adults[16, 17]. Seasonal influenza A and B viruses have a conserved epitope in the NA protein which is necessary for enzymatic function[18, 19]. Antibodies that target this epitope inhibit neuraminidase function and virus replication. Neuraminidase antibodies can be potent and broadly reactive [12, 20]. Anti-NA antibodies increase in titer with age and are capable of recognizing influenza strains isolated in many different influenza seasons [12, 19, 20]. Additionally, a subset of anti-NA antibodies raised in a human infection are broadly cross reactive and protective against influenza A and B virus strains [18, 19].

Neuraminidase antibodies can directly inhibit NA function as well as virus replication. Antibodies that bind neuraminidase can inhibit enzymatic activity, presumable through steric inhibition of substrate accessing the active site [12, 15]. Blocking NA activity prevents the virus from properly budding, leading to virions which aggregate at the cell surface [9, 12, 21]. Furthermore, escape mutants that decrease binding of certain active site targeting anti-NA antibodies incur a significant fitness disadvantage in virus replication and enzymatic activity [19]. This is due to mutating residues critical for the enzymatic function which these broadly reactive antibodies target. These studies indicate the NA protein has a highly conserved and critical epitope which can be targeted by neutralizing antibodies. Targeting the NA protein has recently become one strategy for generating a universal influenza vaccine [15, 17, 20, 22]. As such, a polyclonal antibody response to the NA protein assures inhibition of NA function as well as steric hinderance of the HA protein - effectively neutralizing virus entry and release.

While the HA protein is the immunodominant antigen on the influenza virion, previous studies have shown the function and significance of anti-NA antibodies in vaccination and natural infection [12, 15, 20, 23, 24]. However, this immune pressure can also lead to the selection of viruses that have accumulated mutations in NA protein antigenic sites. NA antigenic drift has been suggested to occur at lower frequency than HA antigenic drift but can have an impact on influenza spread and antibody recognition of NA [25-27].

In 2014-2015 a novel genotype emerged in the human H3N2 influenza viruses. This new genotype encoded an N-linked glycosylation at position 245-247 in the N2 NA protein. This glycosylation is located in close proximity to the NA active site and near a known antigenic site [28]. Using infectious clone technology to assess viral fitness and enzymatic activity, we demonstrate that this NA glycosylation prevents binding of inhibitory antibodies but also reduces NA enzymatic activity and virus fitness in human nasal epithelial cell cultures. The fitness cost of this mutation is therefore balanced by the advantage provided through the escape of preexisting immunity, contributing to viruses with this NA genotype becoming the dominant global H3N2 human virus strain.

## Results

Currently, nearly all circulating human H3N2 viruses have a glycosylation sequence at positions 245-247 in the NA protein. To study the effect that 245 NA glycosylation has on virus replication and enzymatic activity, recombinant viruses were generated which encoded either the 2014/15 N2 NA proteins with (245 NA Gly+) or without (245 NA Gly-) the NA 245 glycosylation and a 2014/2015 HA protein. The remaining six influenza virus segments from A/Victoria/361/2011 (H3N2) were used as the virus genetic backbone. These viruses were first characterized on MDCK-SIAT1 cells, which overexpress the human enzyme CMP-*N*-acetylneuraminate:β-galactoside α-2,6-sialyltransferase producing more cell surface carbohydrates with terminal α-2,6 sialic acid [29]. Both viruses showed similar kinetics of infectious virus production and peak infectious virus amounts after a low MOI infection (Fig 1A). In contrast, infection of hNEC cultures at a low MOI with the 245 NA Gly-virus yielded significantly higher amounts of infectious virus for a prolonged period of time when compared to the 245 NA Gly+ virus (Fig 1B). Plaque appearance, morphology and size was then assessed using MDCK cells. Both viruses produced clear, distinct plaques (Fig 1C) of similar size (Fig 1D). This data indicates that while the 245 NA glycosylation does not impact virus replication on immortalized MDCK-SIAT1 or MDCK cells, it has an adverse effect on virus replication in hNEC cultures.

**Figure 1:**
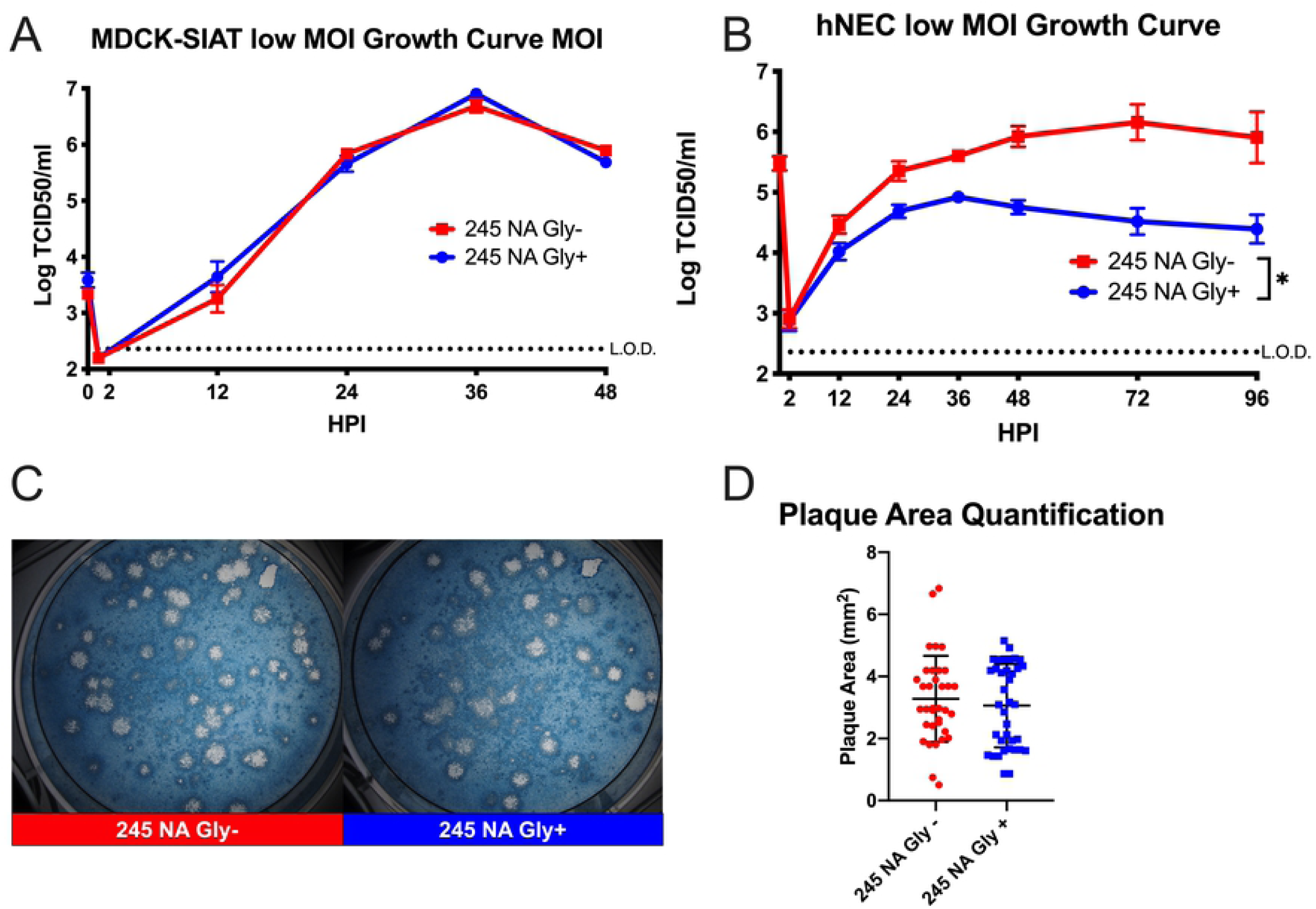
Replication of recombinant H3N2 viruses in MDCK-SIAT1, MDCK or hNEC cultures with or without 245 NA glycosylation. Low MOI growth curves with MDCK-SIAT1 (**A**) or hNEC cultures (**B**) with the indicated recombinant viruses at 32°C. Hours post infection (HPI) on X axis, Log of TCID50/ml on Y axis. Data are pooled from 3 independent experiments with four replicates per virus per experiment (total n = 12 wells per virus timepoint). Data were analyzed with *p<.05 and two-way repeated measures ANOVA with Bonferroni multiple comparison posttest. The limit of detection (L.O.D.) is indicated with a dotted line at log 2.37 TCID50/ml. Error bars in A and B are SEM. (**C**) Plaque assay performed with recombinant 245 NA Gly + and 245 NA Gly - viruses on MDCK cells. (**D**) Quantification of plaque area from 30-50 individual plaques per virus from 3 independent experiments. *p<.05 unpaired T test.

To understand how the addition of a N-linked glycosylation could impact virus replication and protein function, a model of the N2 neuraminidase monomer was generated with UCSF Chimera 3D modeling software. A similar N2 neuraminidase strain (A/Tanzania/2010) was used to highlight key residues and add a complex N-linked glycan at position 245 (Fig 2A) via the online program Glyprot. From the model, it is clear that the 245 N-linked Glycan is uniquely situated near the active site of the protein. To assess whether this N-linked glycan could interfere with the binding of antibodies that target epitopes close to the NA enzymatic active site, the coding sequences for both the 245 NA Gly+ and 245 NA Gly-gene were inserted into a mammalian cDNA expression vector (pCAGGS), with an N-terminal FLAG epitope tag before the stop codon (N-terminus). The NA-FLAG plasmids were transfected into HEK293T cells and the reactivity of the proteins assessed using monoclonal antibodies specific for the NA protein or the FLAG epitope. Three different anti-NA monoclonal antibodies were used. HCA-2 is a rabbit IgG that recognizes a highly conserved 9 amino acid sequence (ILRTQESEC) in the active site of most influenza A and B virus NA proteins. [18, 19]. This antibody was unable to bind to the 245 NA Gly+ protein but showed robust binding of the 245 NA Gly-protein (Fig 2B). The human monoclonal antibodies (235-1C02 and 229-1G02) were also used to study epitope masking. The binding of these antibodies to N2 NA proteins have been described previously [20]. NA proteins encoding amino acid changes at 248 and 429 [20] allow for escape from binding with 235-1C02, suggesting that glycosylation at 245 could inhibit antibody binding to its epitope. In fact, binding of the 235-1C02 to the 245 NA Gly+ protein was not detected but the antibody recognized the 245 NA Gly-protein (Figure 2C). The monoclonal antibody 229-1G03 was previously shown to robustly bind to 245 NA Gly-proteins, but its binding epitope has not been mapped. This antibody can inhibit NA enzymatic activity, suggesting it binds near the NA active site [20]. We found that this antibody recognizes both 245 NA Gly- and 245 NA Gly+ proteins but shows decreased binding to the 245 NA Gly+ protein, suggesting that the 245 NA glycan partially disrupts 229-1G03 antibody epitope recognition (Figure 2D). Taken together these results indicate that the 245 NA glycan masks epitopes in and around the active site of the protein as well as multiple epitopes recognized by human monoclonal antibodies, some of which are potent, broadly reactive inhibitory antibodies. Similar results have recently been reported using the NA protein of the A/Hong Kong/4801/2014 vaccine strain [28].

**Figure 2:**
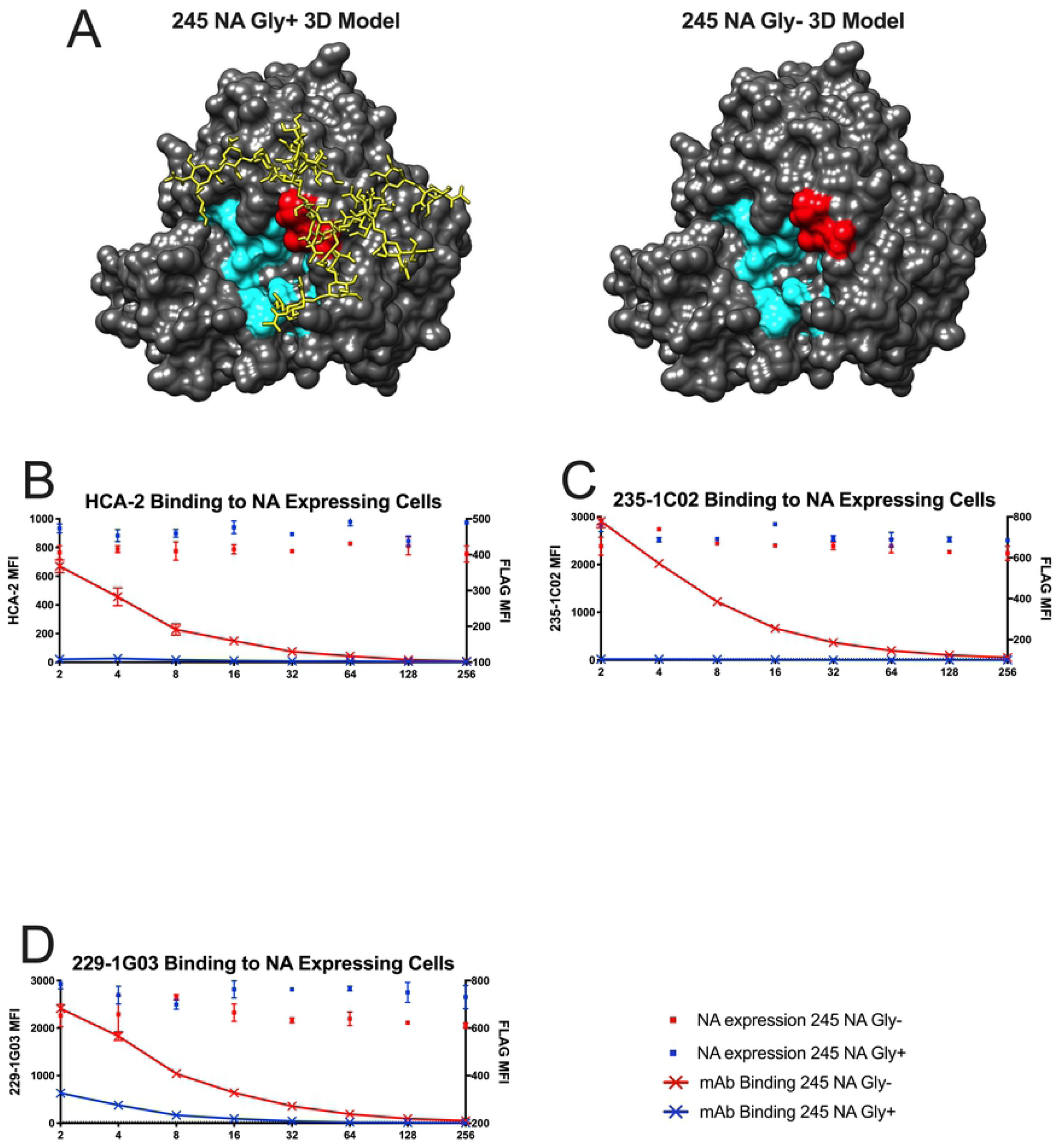
Binding of neuraminidase inhibitory antibodies to cells expressing NA Gly+/- proteins. (**A**) 3D model of N2 NA with (Left, 245 NA Gly+) or without (Right, 245 NA Gly-) the predicted 245 N-glycan. Catalytic and framework residues are highlighted in cyan. Residues 245-247 are highlighted in red. Protein structure modeled and modified via UCSF Chimera, Protein Data Bank ID code 4GZP (Tanzania/2010 N2 NA). A typical complex style N-glycan was added via the Glyprot program. (**B**-**D)** 245 NA Gly+ (blue dots) or 245 NA Gly- (red dots) FLAG-tagged proteins expressed in HEK293T cells. NA expressing cells were incubated with dilutions of monoclonal antibodies HCA2 (**B**), 235-1C02 (**C**) or 229-1G03 (**D**) in addition to a mouse monoclonal antibody recognizing the FLAG epitope (to measure overall NA expression). Red lines indicate mAb binding to cells expressing 245 NA Gly- protein. Blue lines indicate mAb binding to cells expressing 245 NA Gly+ protein. Representative data from 3 experiments. * p <.05 two-way repeated measures ANOVA with Bonferroni multiple comparison posttest.

To understand how 245 NA glycosylation impacted NA function a variety of enzymatic and kinetic activity assays were performed. To standardize NA content, we chose to partially purify virus particles using ultracentrifugation over a sucrose cushion then normalize for NA content using Western blotting with the HCA-2 monoclonal antibody. While the HCA-2 antibody binding to conformationally intact 245 NA Gly+ protein is inhibited, when the protein is denatured, the HCA-2 linear epitope is recognized in both the 245 NA Gly- and Gly+ proteins (Figure 3A) [18, 19]. The NA enzymatic activity was measured using three different NA assays. The Enzyme Linked Lectin Assay (ELLA) uses fetuin (Figure 3B) as a complex carbohydrate substrate which mimics the natural ligands seen by the NA protein during natural infection [30, 31]. The NA-*STAR* (Figure 3C) and NA-Fluor assays (Figure 3D) utilize smaller sialic acid mimics that release luminescent or fluorescent molecules after cleavage. Using all three substrates, the enzymatic activity of 245 NA Gly-was significantly higher than that of the 245 NA Gly+, suggesting that the 245 glycosylation was adversely affecting NA enzymatic activity. This NA activity difference was highest in the ELLA assay, suggesting that the 245 N-linked glycan sterically blocks the full carbohydrate substrate in this assay from the active site. However, the relatively smaller NA-STAR and NA Fluor substrates were still utilized less efficiently by the 245 NA Gly+ protein, suggesting this glycosylation may have more extensive structural effects on the NA active site.

**Figure 3:**
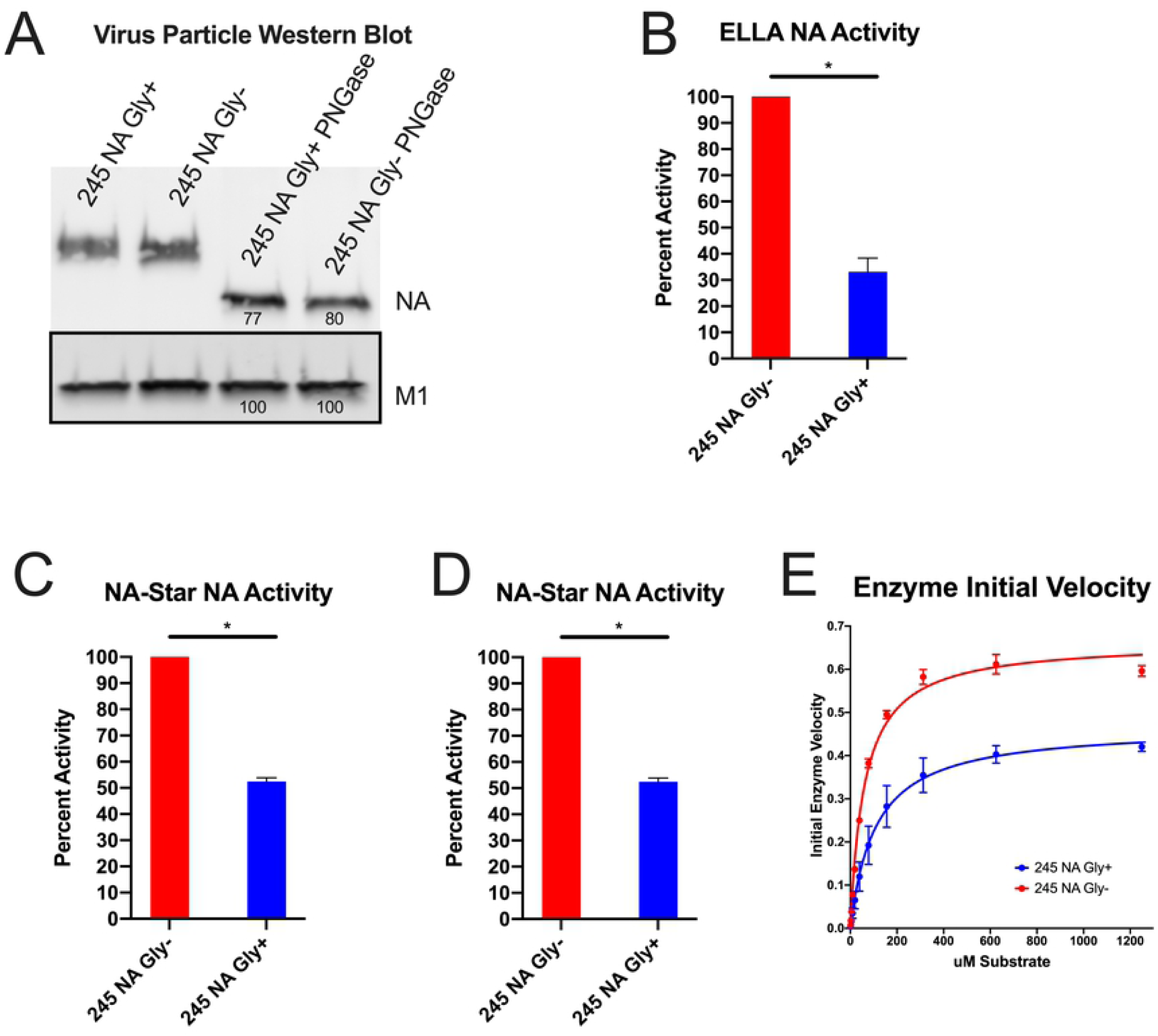
Effect of 245 NA glycosylation on neuraminidase activity. The NA content in partially purified influenza virus particles was measured via SDS-PAGE and western blot (**A**) using HCA-2 mAb to detect NA and M1 antibody GTX125928 to detect M1. Numbers below protein bands indicate measured intensity. NA content was normalized to the M1 content of the same virus sample. With NA content normalized, the NA activity in the partially purified virus preparations was measured in the enzyme linked lectin assay (ELLA) (**B**), NA-STAR assay (**C**) and NA-Fluor MUNANA based assay (**D**). In B, C, and D, 245 NA Gly-enzymatic activity was set to 100. X axis label is viral NA genotype 245 NA Gly+ activity is graphed as a percentage of that activity. (**E**) To assess enzyme kinetics, 245 NA Gly- and 245 NA Gly+ viruses were incubated with a dilution of MUNANA substrate and fluorescence was measured every 60s for 1 hour. Initial velocity plotted as uM product generated per minute. Non-linear regression plotted (line) with individual values (points). * p < .05 unpaired T test. NA and M1 protein content in A were determined using ImageJ software. Enzyme kinetics was determined using a non-linear curve fit Michaelis-Menten equation in Graphpad prism 8.

In addition to bulk activity assays, we performed an enzyme kinetic assay to determine enzyme velocity and affinity for substrate (3E). As expected, the 245 NA Gly+ protein has lower enzyme velocity and a lower affinity for substrate (Fig 3E, Table 1). All of these findings indicate that the 245 NA glycan significantly decreases NA enzymatic activity by decreasing substrate access to the active site of the protein.

**Table 1:** Enzyme kinetics of 245 NA Gly- and 245 NA Gly+ viruses. NA-Flour assay conducted in triplicate, representative of two biological replicates. Values calculated with Graph Pad prism 8 with Michaelis-Menten non-linear regression. 95% confidence interval (CI) shown.

| Test Virus | vMAX (95% CI) | Km (95% CI) | R squared of line |
| --- | --- | --- | --- |
| 245 NA Gly- | .6645 (0.6305 to 0.7001) | 61.55 (50.59 to 74.76) | .9942 |
| 245 NA Gly+ | .4680 (0.4551 to 0.4813) | 108.9 (99.12 to 119.6) | .9988 |

Since the 245 NA glycosylation blocked or decreased binding of the two human monoclonal antibodies 235-1C02 and 229-1G03 and we tested the ability of these antibodies to inhibit viral enzymatic activity. First, viral stocks of 245 NA Gly+ and 245 NA Gly-were equalized via NA content and virus was incubated with a dilution series of the human monoclonal antibodies or oseltamivir. Vehicle (assay buffer) was used for a control and used to subtract background. As expected from the antibody binding studies, the monoclonal antibody 235-1C02 was unable to inhibit the NA enzymatic activity of the 245 NA Gly+ in the NA star assay even at the highest concentration tested (100nm) but inhibited the 245 NA Gly-virus at a concentration of 0.8nm (Figure 4A). The 229-1G03 inhibited both the 245 NA Gly+ and 245 NA Gly-at a concentration of 4.7nm and 1.1nm respectively, suggesting a partial inhibition of inhibitory activity (Figure 4A) via the 245 NA glycan. The same trend is seen in the ELLA assay (4B) with 235-1C02 unable to inhibit the neuraminidase activity of the 245 NA Gly+ virus and 229-G03 showing reduced inhibitory activity. Importantly, in both assays, oseltamivir inhibition was clearly observed and not different between viruses, suggesting that the drug was fully capable of inhibiting NA enzymatic activity irrespective of 245 NA glycosylation status. These results confirm that 245 NA glycosylation can result in reduced inhibitory activity of antibodies that bind near the NA active site. In addition to monoclonal antibody studies we investigated how human convalescent serum from the 2014 through 2016 influenza seasons could inhibit enzymatic activity of the 245 NA Gly+ and 245 NA Gly-protein. We generated H6N2 viruses to avoid the confounding effect that anti-HA antibodies in human serum can have on NA enzymatic activity [30, 32]. Twenty serum samples taken from individuals approximately 28 days after confirmed H3N2 infection were used. Ten serum samples were from patients infected with a 245 NA Gly-virus and 10 from patients infected with a 245 NA Gly+ virus (Table 2). Regardless of the source of serum, the 245 NA Gly+ protein was more resistant to serum based enzymatic inhibition, indicated by a higher concentration of serum needed to inhibit 50% of the enzymatic activity (Fig 4C-E, Table 2) when compared to the 245 NA Gly-virus. In 18 of the 20 serum samples tested, two to three-fold more serum was necessary to inhibit the 245 NA Gly+ protein compared to the 245 NA Gly-protein (Fig 4F). Together these results demonstrate that the 245 NA glycosylation sequence reduces the recognition of serum NA antibodies consistent with antigenic drift of the NA protein.

**Table 2:** Serum samples and 50% NAI values. Serum samples taken from CEIRS study. Serum genotype, 50% NAI (NAI_50_) titer and fold difference shown. Twenty convalescent serum samples taken approximately 28 days after confirmed H3N2 infection used. Ten from 245 NA Gly+ infected patients, 10 from individuals infected with a 245 NA Gly-virus. NAI_50_ values are the highest titer that resulted in at least 50% inhibition of enzyme activity in ELLA assay using H6N2 viruses expressing either 245 NA Gly+ or 245 NA Gly- protein. Data shown from one biological replicate. Each NAI assay was conducted in duplicate and averaged to determine titer.

| Convalescent<br>Serum ID | Serum NA<br>Genotype | 245 NA Gly+<br>NAI<sub>50</sub> Titer | 245 NA Gly-<br>NAI<sub>50</sub> Titer | Fold NA Gly+ /<br>NA Gly- |
| --- | --- | --- | --- | --- |
| 01-23-A-0081 | NA Gly Positive | 80 | 80 | 1 |
| 01-23-A-0023 | NA Gly Positive | 160 | 640 | 4 |
| 01-23-A-0051 | NA Gly Positive | 160 | 320 | 2 |
| 01-11-A-0262 | NA Gly Positive | 1280 | 2560 | 2 |
| 01-21-A-0268 | NA Gly Positive | 1280 | 2560 | 2 |
| 02-11-Pro-0003 | NA Gly Positive | 80 | 80 | 1 |
| 02-11-Pro-0005 | NA Gly Positive | 160 | 320 | 2 |
| 02-11-Pro-0023 | NA Gly Positive | 320 | 1280 | 4 |
| 02-11-Pro-0029 | NA Gly Positive | <40 | 320 | 8 |
| 02-11-Pro-0101 | NA Gly Positive | 160 | 2560 | 8 |
| 01-11-A-0148 | NA Gly Negative | 40 | 80 | 2 |
| 01-11-A-0256 | NA Gly Negative | 1280 | 5120 | 4 |
| 01-11-A-0307 | NA Gly Negative | 640 | 1280 | 2 |
| 02-11-Pro-0006 | NA Gly Negative | 1280 | 1280 | 1 |
| 01-21-A-0192 | NA Gly Negative | 320 | 640 | 2 |
| 02-11-Pro-0030 | NA Gly Negative | <40 | <40 | 1 |
| 02-11-Pro-0036 | NA Gly Negative | <40 | 160 | 4 |
| 02-11-Pro-0056 | NA Gly Negative | 160 | 320 | 2 |
| 02-11-Pro-0057 | NA Gly Negative | 160 | 320 | 2 |
| 01-21-A-0185 | NA Gly Negative | 160 | 320 | 2 |

**Figure 4:**
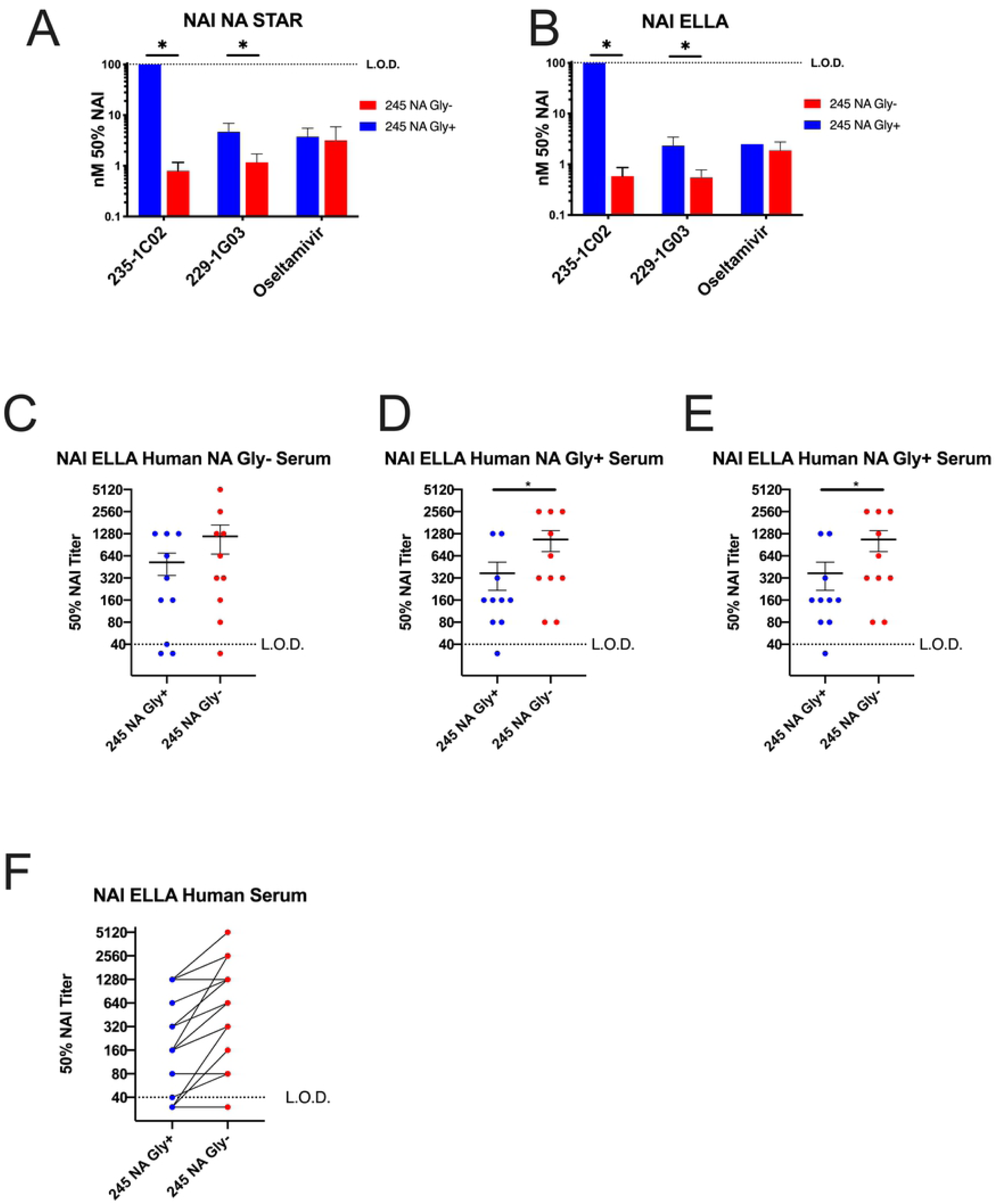
Effect of inhibitory antibodies and human serum on NA enzymatic function ELLA and NA Star. Concentration of N2 monoclonal antibody needed to inhibit 50% of NA activity of 245 NA Gly+ and 245 NA Gly- viruses in NA-STAR (**A**) or ELLA (B) NA activity assays using partially purified H3N2 viruses. Upper limit of detection shown with a dotted line in A and B, indicating the highest concentration of inhibitory antibody used (100nM). (**C-E**) NA inhibition (NAI) ELLA assay performed with human convalescent serum from patients with confirmed H3N2 infection using H6N2 recombinant viruses. Virus content equalized via plaque assay. Convalescent serum NAI assay from all patients with confirmed H3N2 infection (**C**) with NA Gly- virus (**D**), and NA Gly+ virus (**E**). X axis label indicates virus NA genotype. All patient serum samples with connecting lines between matched serum samples (**F**). Serum samples from the same individual are connected to indicate relative activity to the 245 NA Gly+ and 245 NA Gly- viruses. Dotted line shown is lower limit of detection in C-F, highest concentration of convalescent serum used (1:40 dilution). * p<.05 paired T-Test

Neuraminidase inhibitory antibodies have previously been shown to inhibit virus replication by inhibiting enzymatic activity of the protein or by inducing a cellular immune response through antibody dependent cellular cytotoxicity (ADCC) [12, 20, 30, 33] or a combination of both. With two recombinant viruses only differing in the 245 NA glycosylation sequence, we sought to understand how this glycan would impact the ability of 229-1G03 and 235-1C02 to neutralize virus infectivity. Using the two recombinant viruses we found that the antibody 235-1C02 was unable to neutralize the 245 NA Gly+ virus, but effectively neutralized the 245 NA Gly-virus at an average concentration of 1.3nm (Fig 5A). Using 229-1G03, we found this antibody was able to neutralize both 245 NA Gly+ and 245 NA Gly-viruses, with an average concentration of 6.4nm and 1.5nm respectively, indicating somewhat reduced neutralizing activity against the 245 NA Gly+ virus (Fig 5B). Using the experimentally determined 50% neutralizing antibody concentration with the 245 NA Gly-virus in Fig5A and 5B, a multistep growth curve in the presence or absence of these antibodies was performed. Figure 5C demonstrates that the 245 NA Gly+ virus was not impacted with the 235-1C02 antibody, as no significant difference was found in infectious virus production comparing human IgG isotype (clone IGHG1) and 235-1C02. However, antibody 229-1G03 did significantly decrease infectious virus production of the 245 NA Gly+ virus, showing a partial ability to neutralize infectious virus, consistent with the binding (Fig 2) and enzymatic inhibition results (Fig 4). This suggests that the epitope this antibody binds is partially accessible on the 245 NA Gly+ protein. In Figure 5D, both human monoclonal antibodies significantly decreased infectious virus production of the 245 NA Gly-virus to near undetectable levels, suggesting potent neutralizing activity. These results confirm our previous findings with protein binding (Fig 2) and enzymatic inhibition (Fig 3). The 245 NA glycan prevents NA active site-specific antibodies from binding and inhibiting the NA protein, and significantly decreases antibody mediated neutralization of other NA specific neutralizing antibodies.

**Figure 5:**
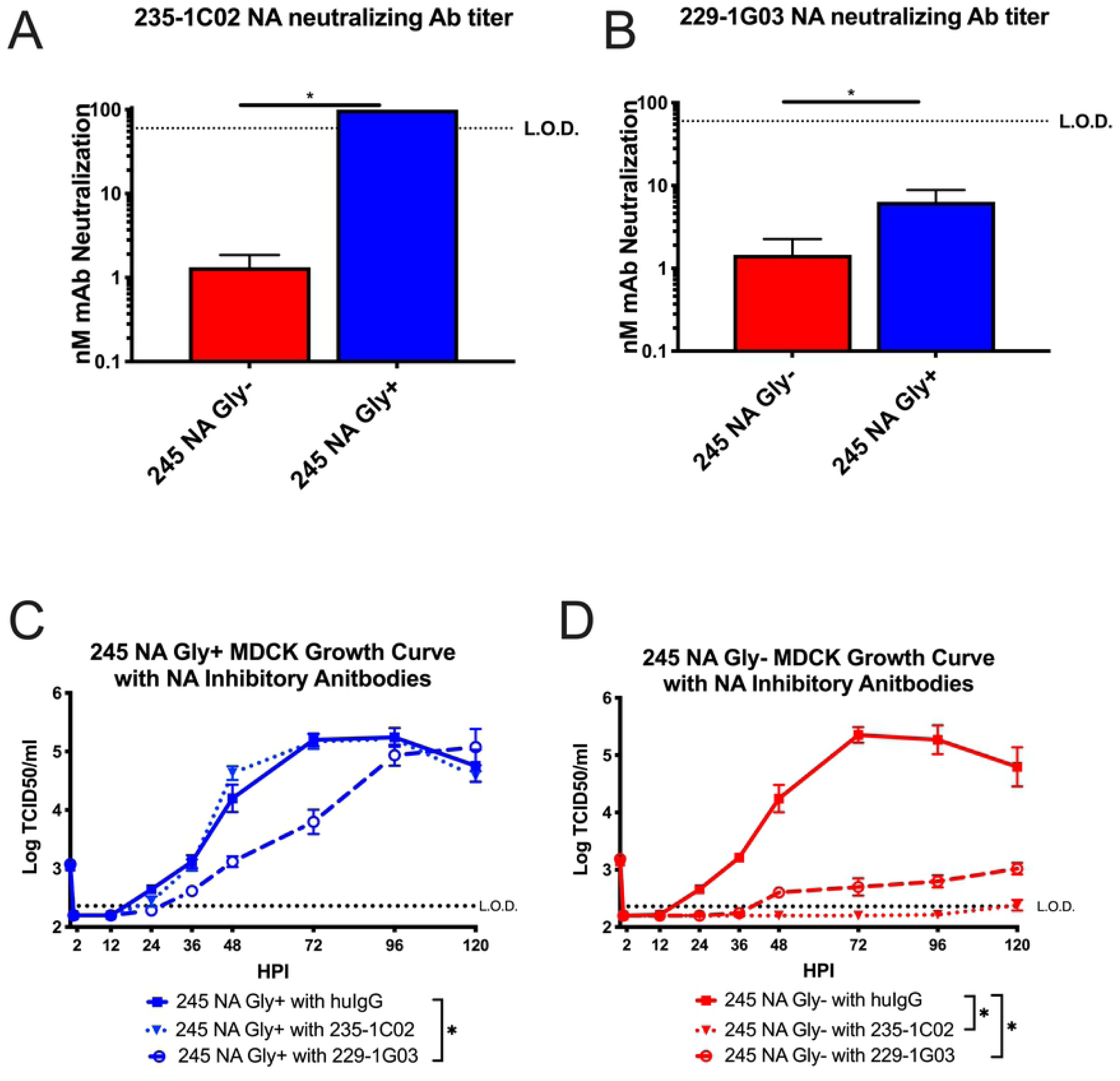
Effect of neuraminidase activity inhibiting antibodies on virus growth. The concentration of anti-neuraminidase monoclonal antibody 235-1C02 (**A**) and 229-1G03 (**B**) needed to neutralize 50% of the infectivity of 245 NA Gly – or 245 NA Gly+ viruses was determined on MDCK cells. Low MOI growth curve with recombinant viruses on MDCK cells (**C-D**). Hours post infection (HPI) on X axis, Log of TCID50/ml on Y axis. MDCK cells were infected with an MOI of .001 with either 245 NA Gly+ virus (**C**) or 245 NA Gly- virus (**D**). After 1hr of inculation, viruses were treated with either human IgG isotype control (clone IGHG1), mAb 235-1C02 or mAb 229-1G03. Dotted line in A and B indicated upper limit of detection, highest concentration of mAb used (100nM). .* p< .05 unpaired T test A and B. Dotted line in C and D indicated lower limit of detection, 2.37 TCID50/ml. Data are pooled from 3 independent experiments with four replicates per virus per experiment (total n = 12 wells per virus timepoint) in C and D Error bars in C and D is SEM. * p<.05 two way repeated measures ANOVA with Bonferroni multiple comparison posttest in C and D.

## Discussion

In this study we demonstrated that the recently acquired 245 N-linked glycosylation site in the NA protein of currently circulating human H3N2 viruses significantly alters the function and antigenicity of the NA protein. The 245 NA glycan decreased in vitro replication on primary hNECs but did not decrease replication on immortalized MDCK cells nor decrease plaque area of isogenic viruses (Fig 1). This suggests that some aspect of primary hNECs, likely the presence of respiratory mucins, is decreasing virus replication. Neuraminidase is necessary for virus motility through mucins [10, 34] and decreasing NA enzymatic activity likely decreases the ability of the virus to move through mucus. The decrease in neuraminidase activity found in three separate activity assays (Fig 3) but was most pronounced when fetuin was used as a substrate, indicating recognition of sialic acid on longer carbohydrate chains is especially affected by 245 NA glycosylation. We conclude that the 245 NA glycan likely blocks substrate access to the active site and decreases enzymatic activity. Decreasing enzymatic activity is likely tied to a decrease in replication seen in mucin secreting hNECs but not seen in immortalized MDCK cells, which to this point have not been shown to secrete mucins.

The presence of a glycosylation site at NA 245 did not affect NA sensitivity to the antiviral drug oseltamivir. While oseltamivir access to the NA active site may be reduced due to the 245 NA glycosylation in a manner similar to that seen with the other enzyme substrates used, subsequent release of oseltamivir is most likely not effected, resulting in efficient inhibition of NA enzymatic activity. Further studies of the kinetics of oseltamivir inhibition of 245 NA Gly+ and 245 NA Gly-viruses could provide additional insights into this observation.

Recently there have been attempts to map the antigenic regions of the NA protein. The 245 NA glycan is located near the enzymatic active site ([28], Fig 2) and is poised to mask this region of the NA protein. We sought to understand how this glycosylation, which incurs a significant fitness disadvantage as judged by virus replication in hNEC cultures, could still fix in the human H3N2 virus population in such a short timeframe. Through multiple assays we found this glycan has an important role in masking NA antigenic sites. This glycan blocks NA active site-specific antibodies from binding (Fig 2), prevents NA active site-specific antibodies from inhibiting enzyme function (fig 4), and blocks the ability of active site antibodies to neutralizing virus replication (fig 5). Additionally, another NA specific monoclonal (229-1G03) antibody with an as yet undefined binding epitope is partially blocked from binding to their epitope by this glycan (Fig 2, 4, 5), suggesting that the 245 NA glycan masks multiple epitopes on the NA protein. This inhibition of NA inhibitory antibody activity was shown with specific monoclonal antibodies and with polyclonal serum from H3N2 infected individuals. The ability to escape from preexisting NA immunity therefore provides a significant fitness advantage for the virus. While we used serum antibody levels to show reduced activity towards 245 NA Gly+ viruses, assessing escape from NA antibody in respiratory tract secretions would be more relevant. This antibody evasion presumably counters the reduced replication of 245 NA Gly+ viruses in hNEC cultures, resulting in a virus whose overall fitness for infecting humans is increased compared to 245 NA Gly-viruses. Since the virus replication fitness deficits were only observed in hNEC cultures while the antibody inhibition of virus replication was evident in immortalized cell lines, our use of hNEC cultures has allowed for a more complete understanding of the effects of 245 NA glycosylation on virus fitness.

Neuraminidase works in conjunction with the HA receptor of the influenza virus to infect and spread virus effectively [35, 36]. As such, studying the NA and HA proteins together is crucial to understanding viral evolution. Both the HA and the NA protein interact with the same ligand, sialic acid, and thus balancing each proteins’ affinity for this ligand is critical to the influenza cycle [35-38]. Both proteins are necessary for in vivo replication, but the nuance of their interaction is important as well. Too strong of an HA-sialic acid interaction compared to NA activity results in the HA protein being trapped in respiratory mucins or not being able to release progeny virions from the infected cell [39, 40]. On the other side of the spectrum, too weak of an HA-sialic acid interaction compared to NA activity results in removal of sialic acid receptors before the HA protein can engage its ligand and initiate infection. This fine balance between affinity for sialic acid impacts virus fitness [36, 37]. Whether the adverse effects of 245 NA Gly+ are observed with more recent H3 HA proteins should be investigated to determine whether HA mutations that compensate for the reduced 245 NA Gly+ enzymatic activity have fixed in human H3N2 viruses.

In recent years the NA protein has had renewed interest as a relatively conserved protein that’s an attractive vaccine target[12, 22, 41]. In some respects, the NA protein is an excellent candidate for a universal vaccine. A single monoclonal antibody can neutralize decades of influenza virus isolates regardless of strain at nanomolar amounts. Neuraminidase inhibitory antibodies can inhibit viral spread, and replication at multiple stages of the virus life cycle [42]. Finally, many different studies show that NA inhibitory antibodies can decrease disease severity, virus transmission or provide sterilizing immunity [5, 14, 18-20, 24].

Antibody response to NA are not induced effectively in all age groups by current influenza vaccines because the amount of NA is not standardized in vaccine preparations and the NA protein conformation is more sensitive to the current vaccine production methods than the HA protein[12, 20, 43, 44]. While other methods for inducing NA immunity are being developed, our data show that two amino acid changes in N2 NA can lead to escape from antibodies that bind to one of the most universal antigenic sites of the protein. It is important that future studies of universal influenza vaccines utilize a multi-epitope vaccine that would require multiple mutations from the virus to escape the vaccine-induced immunity.

This study highlights the necessity to consider multiple aspects of the NA protein in regard to vaccine production and virus evolution. Decades of influenza research have focused on the HA protein for vaccine development, viral evolution and pandemic potential. As the interest in NA protein as a vaccine increases, many of the lessons learned studying influenza HA may also be applied to NA. The NA protein is immunogenic and can provide protection against many strains of influenza viruses[41]. However, like the HA protein, the NA protein can undergo antigenic drift and evade the humoral immune response. As immune pressure mounts due to a renewed vaccination effort at targeting NA protein, the NA protein will likely also become a “moving target” for vaccine development, in a manner similar to what has already been documented for the HA proteins.

## Materials and Methods

### Cell Lines and Primary Cells

Madin-Darby Canine Kidney Cells (MDCK) and human embryonic kidney cells 293T (HEK293T) were maintained in complete medium (CM) consisting of Dulbecco’s Modified Eagle Medium (DMEM) supplemented with 10% fetal bovine serum, 100U/ml penicillin/streptomycin (Life Technologies) and 2mM Glutamax (Gibco). Human nasal epithelial cells (hNEC) were isolated from non-diseased donor tissue following endoscopic sinus surgery. Cells were grown, differentiated and maintained at the air liquid interface as previously described [45]. hNEC differentiation medium and maintenance medium was prepared as previously described [45-47]. hNEC cultures were used for low MOI growth curves only when fully differentiated. All cells were maintained at 37°C in a humidified incubator supplemented with 5% CO_2_.

### Plasmids

The plasmid pHH21 was used to generate full length influenza hemagglutinin (HA) or neuraminidase (NA) plasmids for recombinant virus production. Briefly, viral RNA was isolated from the clade 3c.2a H3N2 viruses A/Bethesda/P0055/2015 (NA Gly+ ID 253812) and A/Columbia/P0041/2014 (NA Gly-ID 253817) with a Qiagen mini-vRNA isolation kit. Gene specific primers with cloning sites for H3N2 neuraminidase or hemagglutinin were used to create cDNA via a one-step RT-PCR reaction (SuperScript III-Platinum Taq mix, ThermoFisher Scientific). The cDNA products were cut with appropriate restriction enzymes, column purified (QIAquick PCR Purification kit) and ligated with restriction enzyme cut-pHH21 using T4-ligase (New England Biolabs, NEB). Ligation products were transformed into DH5a (NEB) cells and colonies were mini-prepped (QIAprep spin mini-prep) and Sanger sequence verified. Sequence verified colonies were maxi-prepped (ZymoPURE) and used for recombinant virus preparation. Since the HA amino acid sequence between A/Bethesda/55/2015 is identical to A/Columbia/41/2014, A/Bethesda/55/2015 HA-pHH21 plasmid was used for both H3N2 viruses. The codon at amino acid position 160 in HA (H3 numbering, Threonine) was modified via site-directed mutagenesis (Agilent) from the wild type (ACA, Thr) to a new codon (ACT, Thr) less likely to revert to a lysine codon-which occurred frequently during previous attempts to virus rescue.

H6 hemagglutinin-pHH21 was synthesized by Genscript (www.genscript.com) in the pHH21 vector. The H6 HA coding sequence from A/Environment/Hubei-Jinzhou/02/2010 [48] was inserted into pHH21 flanked by human H3 5’ (*GCAAAAGCAGGGGATAATTCTATTAACC*) and 3’ (*TAAGAGTGCATTAATTAAAAACACCCTTGTTTCTACTAA*) UTR sequences. After gene synthesis, two mutations (Q223L and G225S) were added in the HA coding sequence to increase HA protein binding to 2,6 sialic acid [49]. The gene product was transformed into DH5a (NEB) and maxi-prepped for recombinant virus production. pHH21 plasmids encoding the internal segments for A/Victoria/361/2011 (H3N2, rVic recombinant viruses) or A/WSN/33 (H6N2, rWSN recombinant viruses) were generated as previous described [50].

The plasmid pCAGGS was used for transient expression of C-terminal flag-tagged NA Gly+ or NA Gly-neuraminidase proteins. C-terminal flag tag (DYKDDDDK) was added to pHH21-NA encoding plasmids via site directed mutagenesis (Agilent). cDNA was generated from the pHH21-NA flag plasmids with Q5 Hot-Start PCR (NEB). This cDNA product was then cloned into the mammalian expression vector pCAGGS for transient transfection experiments as previously described [51].

### Recombinant Virus Production

Recombinant H3N2 or H6N2 viruses were generated using the 12-plasmids reverse genetics system as previously described [50]. Briefly HEK293T cells were plated at 50% confluency 1 day before transfection in complete media. On the day of transfection, media was replaced with serum free Opti-MEM. HEK293Ts were then transfected with eight plasmids encoding full length influenza segments in the pHH21 vector (PB2, PB1, PA, HA, NP, NA, M, NS) and four plasmids encoding the influenza replication proteins in the pcDNA3.1 vector (PB2, PB1, PA and NP). At one day post transfection 5ug/ml N-acetyl trypsin was added to the transfection reaction. MDCK cells were over-laid four hours post trypsin treatment. Every 24 hours post MDCK-overlay virus containing supernatant was sampled for virus production. Fresh Opti-MEM with 5ug/ml N-acetyl trypsin was added when a sample was taken. Virus from the transfected cell supernatants was plaque purified as described below, sequenced, and used to generate seed stocks by infecting MDCK cells at an MOI of 0.001. Working stocks were generated from sequence confirmed seed stocks by infecting MDCK cells at an MOI of .001 as described below.

### Plaque Assay

MDCK cells were grown in complete medium to 100% confluency in 6-well plates. Complete medium was removed, cells were washed twice with PBS containing 2mm calcium magnesium (PBS+) and 400uL of inoculum was added. Cells were incubated at 32°C for 1hour with rocking every 15 minutes. After 1hr, the virus inoculum was removed and phenol-red free DMEM supplemented with3% BSA (Sigma), 100U/ml pen/strep (Life Technologies), 2mM Glutamax (Gibco), 5mM HEPES buffer (Gibco) 5ug/ml N-acetyl trypsin (Sigma) and 1% agarose was added. Cells were incubated at 32°C for 3-5 days and then fixed with 4% formaldehyde. After removing the agarose, cells were stained with napthol-blue black. Plaque size was analyzed in ImageJ [52]. For recombinant virus production, virus plaques were picked with a pipette instead of fixing with formaldehyde and placed in IM and stored at −80°C for later seed stock generation.

### Virus seed and working stocks

For generation of recombinant virus seed stocks, 400ul of plaque picked virus was added to confluent MDCK cells plated in 6 well plates and infected for 1hr as previously described [49, 51]. The plaque pick inoculum was removed and infection media (IM) was added. Infection medium (IM), consisted of DMEM with .3% BSA (Sigma), 100U/ml pen/strep (Life Technologies), 2mM Glutamax (Gibco) and 5ug/ml N-acetyl trypsin((Sigma)). Cells were placed in a 32°C incubator and monitored daily for CPE. Seed stock was harvested between 3 and 5 days or when CPE reached approximately 75-80%. Seed stocks were then sequenced and infectious virus titer determined by TCID50. A working stock for each virus was generated by infecting confluent MDCK cells in a T75 flask at an MOI of .001 for 1 hour at 32°C. The inoculum was removed, and IM was added. Cells were monitored daily for CPE and working stock harvested when CPE reached approximately 75-80%. Working stocks were sequenced verified and infectious virus determined via TCID50 as described below.

### Low-MOI Infections

Low-MOI growth curves were performed at an MOI of .001 in MDCK cells and .01 in hNEC cultures. MDCK cell infections were performed as described above. After the infection, the inoculum was removed and the MDCK cells were washed three times with PBS+. After washing, IM was added and the cells were placed at 32°C. At the indicated times post inoculation, IM was removed from the MDCK cells and frozen at − 80°C. Fresh IM was then added. For low-MOI growth curves in the presence of monoclonal antibodies, the indicated antibodies were added to the IM after the virus was allowed to attach to cells. In low-MOI hNEC growth curves, the apical surface was washed three times with PBS and the basolateral media was changed at time of infection. hNEC cultures were inoculated at an MOI of .01. hNEC cultures were then placed in a 32°C incubator for 2 hours. After inoculation, the hNECs were washed three times with PBS. At the indicated times, 100ul of IM without N-acetyl trypsin was added to the apical surface of the hNECs. The hNECs were then incubated for 5 minutes at 32°C and the IM was harvested and frozen at −80°C. Basolateral media was changed every 48hrs post infection for the duration of the experiment.

### TCID50

MDCK cells were seeded in a 96 well plate 2 days before assay and grown to 100% confluence. Cells were washed twice with PBS+ then 180uL of IM was added to each well. Ten-fold serial dilutions of virus was created and then 20uL of the virus dilution was added to the MDCK cells. Cells were incubated for 6 days at 32°C then fixed with 2% formaldehyde. After fixing, cells were stained with napthol blue-black, washed and virus titer was calculated[49, 51].

### Transient Transfection for NA-Flag expressing cells

Transient transfection of HEK293T was performed with TransIT-LT1 per the manufacturers protocol. Briefly, cells were grown in complete medium until time of transfection to roughly 50% confluency. On the day of transfection, complete medium was removed and replaced with Opti-MEM serum free medium. Opti-MEM, TransIT-LT1 and 2.5ug of plasmids encoding gene of interest were mixed then added to HEK293T cells. At 16hr post transfection wells were used for flow cytometric analysis.

### NA Antibodies

NA specific monoclonal antibodies 229-1G03, 235-1C02 and HCA2 were used to assess binding to NA proteins. 229-1G03 and 235-1C02 were provided by Patrick Wilson [20]. 235-1C02 binds to residues 249 and 428 on the NA protein as described and the 229-1G03 binds to an as yet uncharacterized epitope on the N2 NA protein. HCA-2 monoclonal antibody was provided by Sean Li [18]. HCA-2 binds to a known, highly conserved epitope in the active site of the NA protein, residues 222-230 ILRTQESEC. To assess antibody binding to expressed NA proteins, all monoclonal antibodies were diluted to 1ug/ml 1X PBS (Quality Biologics) containing .1% BSA, (Sigma) was used throughout antibody staining protocol (FACS buffer). The antibodies were then serially diluted 1:2 in FACS buffer. Mouse anti-FLAG (clone M2, Sigma) was diluted in FACS buffer to 1ug/ml. For western blotting mouse anti-FLAG and anti-influenza M1 antibody were diluted to 2ug/ml in blocking buffer. Antibodies were diluted in IM for virus neutralization assays. For low MOI growth curve viral inhibition, NA inhibitory antibodies were diluted in IM + 5ug N-Acetyl Trypsin. 229-1G03 was diluted to 1.5nm, 235-1C02 1.3nm, and human IgG isotype clone IGHG1 diluted to 5nM.

### Secondary Antibodies

Secondary antibodies were used to detect binding of primary unconjugated monoclonal antibodies. Goat anti-Mouse IgG Alexa Fluor 488, Goat anti-Rabbit IgG Alexa Fluor 647 and Goat anti-Human IgG Alexa Fluor 647 were used at 1ug/ml concentration in FACS buffer (ThermoFisher Scientific). For western blotting, all secondary antibodies were diluted in blocking buffer at a concentration of 1ug/ml.

### Human Serum and Ethics statement

Convalescent human serum obtained through the JH-CEIRS study (HHSN272201400007C) were used in this study. Serum samples were treated with receptor destroying enzyme (Cosmos Biological) and heat treated according to the manufacturer’s protocol for use in ELLA studies. The Institutional Review Board at the Johns Hopkins University School of Medicine provided ethical approval of the study (IRB00052743). Patients were approached by trained clinical coordinators who obtained written, informed consent before collecting specimens, demographic and clinical data using a standard questionnaire. Data was confirmed by examination of the patient’s electronic health record. All data was de-identified.

### Flow Cytometry

HEK293T cells were detached with .05% Trypsin-EDTA (Life Technologies) and fixed with 2% paraformaldehyde (Affymetrix) at room temperature for 15 minutes. Cells were washed with FACS buffer after fixation and stained with the indicated amounts of human or rabbit monoclonal antibody and anti-FLAG mouse monoclonal antibody. Cells were washed twice in FACS butter between each antibody incubation step. Cells were analyzed on a BD-FACS Calibur and data analyzed with FlowJo V10.5.3 software. Geometric mean was used to identify mean fluorescence intensity (MFI).

### Partially Purifying Virus Particles

Virus partially purified by ultracentrifugation over a sucrose cushion for SDS-PAGE and western blotting. Clarified virus working stock supernatant was overlaid onto a 25% sucrose-NTE (100nM NaCl (ThermoFisher Scientific), 10mM Tris (Promega) and 1mM EDTA (Sigma)) buffer. Virus was centrifuged at 27,000 RPM in a SW-28 rotor in a Beckman Coulter Optima L90-K UltraCentrifuge for 2 hours. After the first ultracentrifugation, the supernatant was removed. The virus pellet was re-suspended in PBS. Pellet was further concentrated by ultracentrifugation in an SW-28ti rotor at 23,000 RPM for 1hr. The pellet was resuspended in PBS for use in NA activity, western blotting and PNGase assays.

### PNGase, SDS-PAGE and Western Blotting

Partially purified virus particles were used for SDS-PAGE. For PNGase treatment, the PNGase kit from (NEB) was used per manufacturer’s instructions. After PNGase treatment, all samples were treated with 4X-Laemli buffer (Bio-Rad) containing 250mM dithiothreitol (DTT, ThermoFisher Scientific) and boiled at 100°C for 5 minutes. Samples were run on 4-20% Mini-PROTEAN TGX gels (Bio-Rad) with an All-Blue precision plus protein ladder (Bio-Rad) at 70v. Proteins were transferred onto an immobilon-FL membrane (Millipore) at 75v for 1hr. After transfer, membranes were blocked with blocking buffer (PBS containing .05% Tween-20 (Sigma) and 5% non-fat milk (Bio-Rad)) for 1 hour at room temp. Primary antibody (HCA2 and anti-M1) was incubated overnight at 4°C in blocking buffer. Membranes were washed in PBS with .05% Tween-20 (wash buffer). Secondary antibody was added for 1hr at room temperature (25°C) in blocking buffer then washed again in wash buffer. Blots were imaged and analyzed with the FluorChem Q system (Proteinsimple).

### NA-*Star* Assay

NA-Star Influenza Neuraminidase Inhibitor Resistance Detection Kit assay was performed according to manufactures specifications (ThermoFisher Scientific). Briefly, serial two fold dilutions of human serum or monoclonal antibodies were mixed in NA-STAR assay buffer. An equal volume of partially purified virus diluted in NA-Star assay buffer was added to the antibody dilutions. This mixture of virus and antibody was placed in a 96 well white opaque plate and incubated at room temp for 30 minutes with gently horizontal shaking. After incubation, 10ul of 1X NA-*Star* substrate was added and the plates were incubated at room temp for an additional 30 minutes while shaking. After adding substrate, accelerator was added and plates were read immediately by measuring luminescence on a FilterMax F5 multimode microplate reader. To assess overall NA activity, no monoclonal antibody was added. Data was analyzed in Prism (GraphPad) and 50% inhibition was defined as antibody or serum concentration that resulted in at least 50% inhibition of NA activity compared to virus without antibody.

### Enzyme Linked Lectin Assay

Enzyme linked lectin assays (ELLA) were performed as previously [30, 31]. Flat-Bottom Nunc MaxiSorp plates (ThermoFisher Scientific) were coated with 100ul of fetuin (Sigma) at 25ug/ml. Plates were kept at 4°C for at least 18 hours, up to 1 month before use. Monoclonal antibodies, human serum or oseltamivir were serially two-fold diluted in Dulbecco’s phosphate buffered saline with calcium and magnesium (ThermoFisher Scientific) containing 1% BSA (Sigma) and .2% Tween-20 (referred to as sample buffer). Dilutions were performed in 60ul in duplicate on a Nunclon Delta Surface Round bottom 96 well plate. Virus was added to sample buffer, and 60ul of virus was added to the dilution plate. For monoclonal antibody and inhibitor experiments recombinant H3N2 virus was used. For human serum, recombinant H6N2 virus was used. NA content was equalized via western blotting for H3N2 or virus content equalized via plaque assay for H6N2. Fetuin coated plates were washed immediately before addition of 100ul virus premixed with antibody, serum or oseltamivir. Plates were covered with a plastic lid then placed in 37°C incubator with 5% C0_2_ for 16-18 hours overnight. The following day, plates were washed six times with PBS containing .05% Tween 20 (referred to as PBST). After the last wash, 100ul of biotinylated peanut agglutinin lectin at 1ug/ml was added to every well and incubated at room temperature for 2 hours. After peanut lectin addition, plates were washed three times with PBST. Next, 100ul of 1ug/ml streptavidin-horse radish peroxidase (Millipore Sigma) was added to every well and plates were incubated at room temperature for 1 hour. Plates were then washed 3 times with PBST before the addition of 100ul of .5mg/ml *o*-Phenylenediamine (Sigma) diluted in phosphate-citrate buffer with sodium perborate (Sigma). Plates were incubated for 10 minutes at room temperature and reactions were stopped and developed by addition of 100ul of 2N sulfuric acid diluted in water. Absorbance was read at 405nm on a FilterMax F5 multimode microplate reader (Molecular Devices). To assess NA activity, no monoclonal antibody was added. Data was analyzed in Prism (GraphPad8) and 50% inhibition was defined as antibody or serum concentration that resulted in at least 50% inhibition of NA activity compared to virus without antibody.

### NA-Fluor Assay

NA-Fluor Influenza Neuraminidase Assay was performed according to manufacturer’s specifications and enzyme kinetics experiments performed as previously reported [53]. For enzyme kinetics, MUNANA substrate was serially two-fold diluted in assay buffer on an opaque black 96 well plate. Virus was prepared in assay buffer then added to the plate containing MUNANA substrate dilutions. Fluorescence was measured every 60s for 1 hour after addition of virus on a FilterMax F5 multimode microplate reader (Molecular Devices). Enzyme Vmax and Km was calculated using Prism software (GraphPad).

### NA Neutralizing Antibody Assay

To assess the ability of monoclonal antibodies ability to inhibit virus replication, a neutralizing assay was performed. MDCK cells were plated to 100% confluency on 96 well plates and washed twice with PBS+. A two-fold serial dilution of monoclonal antibody was made in IM + 5ug/ml N-acetyl trypsin at a starting concentration of 100nm in a volume of 60ul in duplicate on round bottom Nunclon plates. Next, 60ul (total of 2,000 PFU) of either 245 NA Gly+ or 245 NA Gly-H3N2 recombinant virus diluted in IM with 5ug/ml N-Acetyl Trypsin was added to the dilution plate and 100ul of the mixture of virus and antibody was then added to MDCK plates. After 6 days plates were fixed with 4% formaldehyde and stained with napthol blue-black as described above. Wells were considered negative for virus replication if the entire monolayer was intact.

### NA Neutralizing Antibody Virus Replication Assay

To study monoclonal antibody inhibition of multistep viral growth, viral replication assays were conducted in the presence of NA monoclonal antibodies or human IgG isotype. Confluent MDCKs were infected with an MOI of .001 as described above. After infection, viral inoculum was removed, the cells washed twice with PBS+ and monoclonal antibodies (235-1C02, 229-1G03 or human IgG isotype clone IGHG1) were added at the indicated concentration in IM containing 5ug/ml n-Acetyl trypsin. Infected cells were incubated at 32°C. At each timepoint post infection, supernatant was removed and stored at −80C. Fresh IM with 5ug/ml N-acetyl trypsin and the indicated antibody was added. Viral titer was determined via TCID_50_.

## Acknowledgements

We thank the members of the Pekosz laboratory, the Klein laboratory, and the Davis Lab for data discussion and feedback. We would also like to thank The Johns Hopkins Department of Emergency Medicine, the Johns Hopkins Department of Infectious Diseases, the Johns Hopkins Applied Physics Lab, Dr. Andrew Lane, Dr. Xuguang (Sean) Li, and Patrick Wilson for providing reagents and cells. The work was supported by CEIRS HHSN272201400007C and T32 AI007417 (HP).

